# Quantifying Viscous Damping and Stiffness in Parkinsonism Using Data-Driven Model Estimation

**DOI:** 10.1101/2021.02.17.431730

**Authors:** Alec Werning, Daniel Umbarila, Maxwell Fite, Sinta Fergus, Jianyu Zhang, Gregory F. Molnar, Luke A. Johnson, Jing Wang, Jerrold L. Vitek, David Escobar Sanabria

## Abstract

Rigidity of upper and lower limbs in Parkinson’s disease (PD) is typically assessed via a clinical rating scale that is subject to biases inherent to human perception. Methodologies and systems to systematically quantify changes in rigidity associated with angular position (stiffness) or velocity (viscous damping) are needed to enhance our understanding of PD pathophysiology and objectively assess therapies. We developed a robotic system and a model-based approach to estimate viscous damping and stiffness of the elbow. Our methodology enables the subject to freely rotate the elbow, while torque perturbations tailored to identify the arm dynamics are delivered. The viscosity and stiffness are calculated based on the experimental data using least-squares optimization. We validated our technique using computer simulations of the arm dynamics and experiments with a nonhuman animal model of PD in the presence and absence of deep brain stimulation (DBS) therapy. Computer simulations of the arm demonstrate that the proposed approach can accurately estimate viscous damping and stiffness. The experimental data show that both viscosity and stiffness significantly decreased when DBS was delivered. Computer simulations and experiments showed that stiffness and viscosity measurements obtained with the proposed methodology could better differentiate changes in rigidity than scores previously used for research, including the work score, impulse score, and modified Unified Parkinson’s Disease Rating Scale (mUPDRS). The proposed estimation method is suitable to quantify the effect of therapies on viscous damping and stiffness and study the pathophysiological mechanisms underlying rigidity in PD.

## 1. Introduction

Parkinson’s disease (PD) is a disorder of the central nervous system that alters movement control and can lead to limb rigidity, tremors, and balance difficulties. Rigidity is a cardinal motor sign of PD that affects 90-99% of patients [1] and is characterized by stiff and inflexible muscles that prevent relaxation. Objectively quantifying rigidity in PD is vital to evaluating therapies and advancing treatments such as deep brain stimulation (DBS). In both clinical and research settings, the Unified Parkinson’s Disease Rating Scale (UPDRS-III) is the most widely used approach for assessing PD motor signs. To obtain the rigidity UPDRS-III sub-score, the examiner manipulates major joints and assigns a 0-4 rating based on the rigidity’s severity[2]. The UPDRS-III scale has low sensitivity to quantify mild severity or small changes in motor signs due to its coarse rating and dependence on human perception, and can exhibit limited inter-rated reliability depending on the rater’s training [3, 4, 5].

Devices and approaches based on force transducers have been developed to objectively and precisely assess rigidity in both human and nonhuman primate models of PD [6, 7, 8]. For example, the system developed by Fung et al. [8] moved the subject’s wrist manually (0.5-1.0 cycles/s) and measured the reaction torque using a torque transducer. The device developed by Mera et al. [7] moved the subject’s elbow with a triangular angular position trajectory (0.5 cycles/s) generated by a motor and measured the reaction torque using a torque transducer. These studies quantified rigidity through the angular impulse and work scores, which are scalar measures based on torque and position measurements resulting from movement trajectories at one single frequency (number of cycles/sec). Impulse and work scores provide a more objective assessment of rigidity than the UPDRS-III scale given that unbiased sensors and actuators are employed for their calculation; however, these scores cannot identify whether rigidity measurements are associated with changes in angular position (stiffness) or velocity (viscous damping).

While the rigidity of upper and lower joints in PD has long been thought to be dependent on the angular position (stiffness), recent research has shown that angular velocity (viscous damping) has a significant impact on rigidity [9]. A study by Park et al. [10] with PD patients and control subjects, in which damping and stiffness were estimated, found that viscosity and stiffness associated with flexion-extension of the wrist correlated with clinical rigidity assessments. The authors in [10] estimated the viscosity and stiffness from the individual flexion and extension torque measured in reaction to movement manually applied by an examiner at 0.2-1.0 cycles/s. They could differentiate PD subjects from controls with their model based on viscosity estimates alone. They also suggested that enhanced long-latency stretch reflexes mediated by the sensorimotor cortices may contribute to the velocity-dependent muscle activation associated with viscosity, whereas background muscle activation and passive resistance may contribute to the stiffness. The work by Park et al. [10] highlights the importance of accounting for both velocity (viscosity) and displacement (stiffness) effects on rigidity measurements.

We built upon the work by Park et al. [10] and Mera et al. [7] to develop a robotic system and rigidity assessment methodology for elbow motion that enables controlled generation of torque perturbations and free motion. The system’s free-motion capability allows us to generate torque perturbations while the subject moves the arm freely, and to separate the effect of our known torque perturbations from the subject’s voluntary torque applied to the robot (unknown disturbances). This capability is instrumental when conducting experiments with animal models of PD and human patients with tremor and dyskinesia. The known torque perturbations were designed to have a frequency content that excites the rotational arm dynamics in order to improve model accuracy. We modeled the rotational arm dynamics with a second-order differential equation. The stiffness and viscosity parameters of the model were estimated using least squares estimation based on the known torque perturbations and measurements of position, velocity, and reaction moment. To understand the benefits and drawbacks of our rigidity assessment methodology compared to rigidity scores typically used for research, we employed both computer simulations and recordings from a nonhuman primate model of PD in the presence and absence of DBS therapy. Computer simulations of the arm and robot rotational dynamics indicate that our estimation approach can accurately identify changes in both viscosity and stiffness. These simulations also show that the work and angular impulse scores capture a combination of the viscosity and stiffness components in a single scalar measurement, but under certain conditions, the work score does not reflect overall changes in rigidity. The animal PD model experiments showed that changes in rigidity associated with DBS could be detected with the mUPDRS (modified UPDRS-III) scores blinded to the examiner, work and angular impulse scores, and data-driven parameter estimation technique. Our estimation technique further revealed that DBS reduced both viscosity and stiffness in the studied subject. Moreover, the differences in viscosity and stiffness detected with our parameter estimation method had a greater effect size than the effect sizes observed with the mUPDRS, work, and angular impulse scores. This study suggests that the proposed parameter estimation approach can be utilized to effectively quantify and characterize viscous and elastic rigidity in parkinsonian subjects, critical to objectively assessing the effectiveness of therapies in both research and clinical settings.

## 2. Methods

### 2.1. Subject

All procedures were approved by the University of Minnesota Institutional Animal Care and Use Committee (IACUC) and complied with United States Public Health Service policy on the humane care and use of laboratory animals. A rhesus macaque rendered parkinsonian via administration of the 1-methyl 4-phenyl 1,2,3,6-tetrahydropyridine (MPTP) neurotoxin [11] was used to evaluate the rigidity assessment methods studied here. We assessed the rigidity of the subject’s left elbow during two conditions: no treatment (OFF) and high-frequency DBS of the right subthalamic nucleus. DBS was delivered to the subject at 130 Hz frequency, 320 *µ*A amplitude, and 60 *µ*s pulse-width. We used a bipolar electrode configuration, in which the anode and cathode of the current source were connected to adjacent electrodes in the DBS array. Rigidity assessments were performed 15 minutes after DBS was turned on to account for possible wash-in effects of DBS on rigidity.

### 2.2. Rigidity Assessment via Rating Scale

Elbow rigidity was evaluated by three blinded examiners with mUPDRS scores ranging from 0 to 3, with increments of 0.5 and 3.0 representing the highest severity. [12].

### 2.3. Experimental Setup

Elbow rigidity was assessed using a robotic system attached to the subject’s forearm, with the axis of the robot’s actuator aligned with the subject’s elbow. The position and velocity of the robot’s arm were controlled using a high torque, AC servo-actuator (39 Nm FHA-C series, Harmonic Drive, US). A force/torque sensor (F/T Sensor: Gamma, ATI, US) measured the torque produced by the subject at the shaft of the actuator. Figure 1 illustrates how the robot is configured and attached to the subject’s arm. We chose to move the elbow along an axis parallel to the subject’s torso to avoid gravity’s effect on the torque measurements.

**Figure 1:**
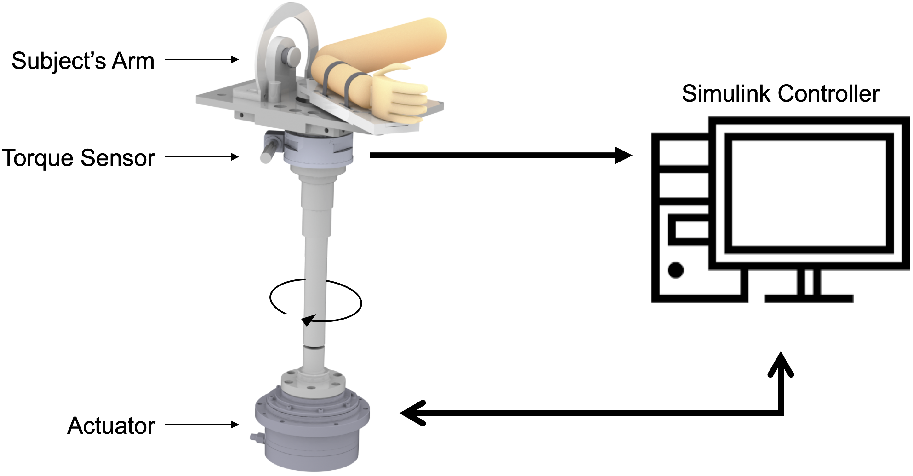
Schematic of subject’s arm with robotic system. The robot was used to perturb the subject’s arm using a high-torque motor and measure the reaction torque applied by the arm.

The rigidity quantification methods studied here were implemented on the robot by using a custom software developed in Simulink Desktop Real-Time (Mathworks, 2016). Custom drivers were used to interface with the actuator and torque sensor. Data streams from the torque sensor were acquired at 100 samples/s. The actuator controller (REL Servo Drive, Harmonic Drive) measured the angular position and velocity of the shaft using an optical encoder with a resolution of 2.25 *·* 10^*−*4^ deg per pulse and transmitted these data to the Simulink Real-Time computer with a sampling rate of 50 samples/s. Physical and software limits were implemented to ensure that the robot’s speed and position did not exceed safety limits.

### 2.4. Rigidity Assessment using Angular Impulse and Work Scores

Sinusoidal position trajectories were applied to the elbow joint at 1.0 Hz with a +/-20 deg range to mimic the clinical UPDRS assessment motion as suggested by Fung et al. [8]. We measured 20 s epochs consisting of the torque produced by the subject’s arm in response to 10 s of movement trajectories (i.e., perturbations). After each epoch, the subject was randomly given a period of 10 or 20 s when the robot’s position was set constant to let the subject rest and minimize the subject’s anticipation of movement. Each experiment lasted around 10 min, with 15 sinusoidal perturbation epochs recorded in each condition (OFF and DBS).

Impulse and work scores were calculated based on the position and torque data acquired during the experiments described above. The first and last second of each epoch were discarded to remove transient effects observed in torque measurements. We used a digital band-pass filter (0.2-5.0 Hz) to remove noise and low-frequency measurements associated with the subject’s active resistance.

The angular impulse score holistically quantifies the inertial, viscous, and elastic forces generated at the elbow. Measurements of this score were computed by rectifying the measured torque and integrating the rectified torque with respect to time for each sinusoidal perturbation. We calculated the angular impulse score over an epoch in which the perturbation frequency was constant. The angular impulse score is given by the following expression

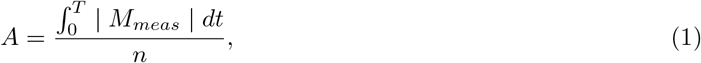

where *M*_*meas*_ is the measured torque (*N · m*), *t* is the time (s), *T* is the integration interval, and *n* is the number of cycles integrated (8 cycles in each epoch). Therefore, the angular impulse score is the average angular impulse over an epoch in which the perturbation frequency is constant.

The work score quantifies the relationship between elbow force and the angular displacement of the elbow. This score represents the mechanical work produced by the elbow during the perturbations. The work score is given by

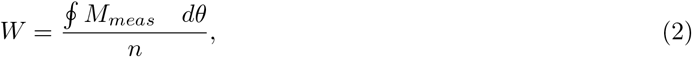

where *M*_*meas*_ is the measured torque (*N · m*), *θ* is the joint angle (rad), and *n* is the number of cycles integrated (8 cycles in each epoch). The net work of each perturbation cycle is defined as the area within the hysteresis loop produced by the torque response with respect to the angular displacements.

### 2.5. Estimation of Viscous and Elastic Rigidity using Admittance Control

#### 2.5.1. Mathematical Abstraction of Elbow Rigidity

A second order, linear, time-invariant differential equation was used to characterize the motion of the arm and associated rigidity about the elbow’s axis of rotation. This equation is given by

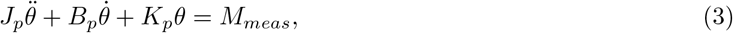

where *J*_*p*_ is the moment of inertia about the elbow’s axis of rotation, *B*_*p*_ is the viscosity, *K*_*p*_ is the stiffness, *θ* is the angular position, and *M*_*meas*_ is the torque measurement. 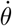 and 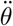 are the first and second temporal derivatives of the angular position (i.e. angular velocity and acceleration). The goal of our estimation approach is to calculate parameters *J*_*p*_, *B*_*p*_, and *K*_*p*_ by using measurements of *θ*, 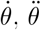,and *M*_*meas*_.

#### 2.5.2. Admittance Controller

Enabling subjects to freely move their elbow during rigidity assessments can help relax their arm, create a more natural environment than that created when the position of the arm is fixed or driven by a position tracking command, and potentially minimize the recurrence of volitional movement. To create a feeling of free motion, one needs to overcome the high torque generated by the actuator’s mechanical gear. We addressed this problem by actively controlling the actuator velocity based on the torque applied by the subject to the shaft. In the robotics field, this strategy is referred to as admittance control [13].

The admittance controller used torque measurements to generate velocity commands that were executed in real-time by the robot actuator. Sensor measurements were filtered before entering the controller using a low-pass filter with a cutoff frequency of 20 Hz. This filter minimizes the propagation of high-frequency noise from the torque transducer to the robot trajectories. The controller’s transfer function emulated the dynamics of a rotational system with damping (*B*_*c*_) and inertia (*J*_*c*_). Figure 2A illustrates how the velocity commands are generated from a given measured torque (subject instruction). Figure 2C shows the block diagram of the velocity generation within the admittance controller. Without the inertial and damping effects in the controller, the system would move the robot’s arm with an unbounded velocity and position when a constant force is applied, resulting in abrupt motion of the arm followed by a sudden stop when the actuator reaches the position limits.

**Figure 2:**
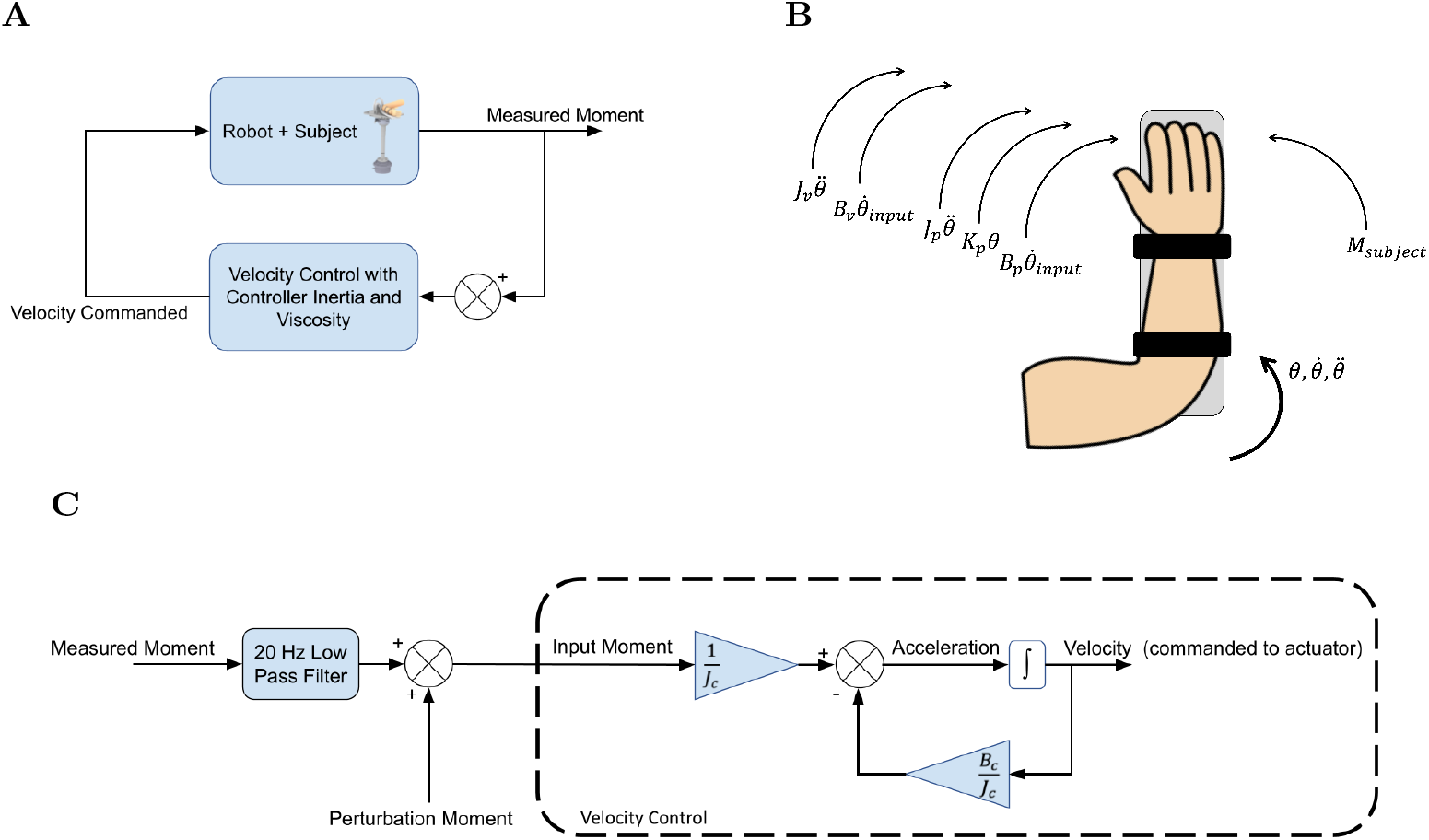
**A:** Block diagram illustrating the implementation of the admittance controller. Torque measurements are fed back to generate the velocity commanded to the actuator. **B:** Free body diagram illustrating the rotational dynamics of the robot, arm, and admittance controller **C:** Block diagram of the admittance controller illustrating how the controller inertia and damping alter the input velocity to reproduce the free motion dynamics based on the subject’s measured torque. A torque perturbation is injected in software is used to excite the arm rotational dynamics and identify the viscous damping and stiffness

Figure 2B shows the physical and computer-generated torque components involved in the arm motion, including the controller’s viscous damping and inertial contributions. We heuristically tuned the controller inertia and viscous damping with the subject’s arm placed on the robot to achieve smooth and low-friction motion when the subject applied a force. The final values were set to *J*_*c*_ = 0.0008*Kgm*^2^*/deg* and *B*_*c*_ = 0.028*Nms/deg*. The torque contributions of the controller inertia and viscosity to the system dynamics are presented in equation 4:

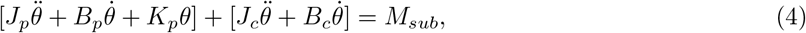

where *M*_*sub*_ is the torque voluntarily applied by the subject. Equation 5 illustrates how the controller and arm-robot dynamics relate to the measured torque (*M*_*meas*_), which in turn can be approximated as the torque generated by the robot actuator (*M*_*act*_).

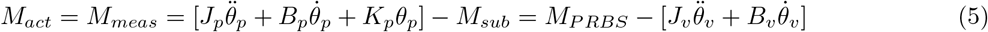

The term *M*_*PRBS*_ represents torque perturbations (pseudo-random binary sequence, PRBS) injected into the torque signal to estimate the effect of these known perturbations on the arm rotational dynamics (i.e., viscosity and stiffness). The PRBS perturbation is described below. One should note that the torque voluntarily applied by the subject (*M*_*sub*_) can be approximated via Equation 5 when estimates of *J*_*p*_, *B*_*p*_, and *K*_*p*_ are available.

#### 2.5.3. Perturbation Design for Parameter Estimation

We used a known perturbation to the torque signal entering the admittance controller to excite the arm dynamics and estimate the rigidity components based on its response to the perturbation. Using both the measured torque and perturbation signal to generate the arm’s motion, we enable the subject to move the arm while perturbing the system dynamics for parameter estimation. We designed the input perturbation torque to excite the subject’s arm at frequencies throughout the expected bandwidth of the arm motion. We estimated the bandwidth of the admittance controller dynamics together with the subject’s arm to have a maximum (worst-case) cutoff frequency of 3.5 Hz. This cutoff frequency was estimated using a second-order model of the arm, based on available stiffness and viscosity data from the human arm, and considering a 50% error in the parameters. A more detailed description of the approach to determine this cut off frequency is outlined in Appendix A. A Bode plot illustrating the frequency response of the arm model is presented in Figure 3A.

**Figure 3:**
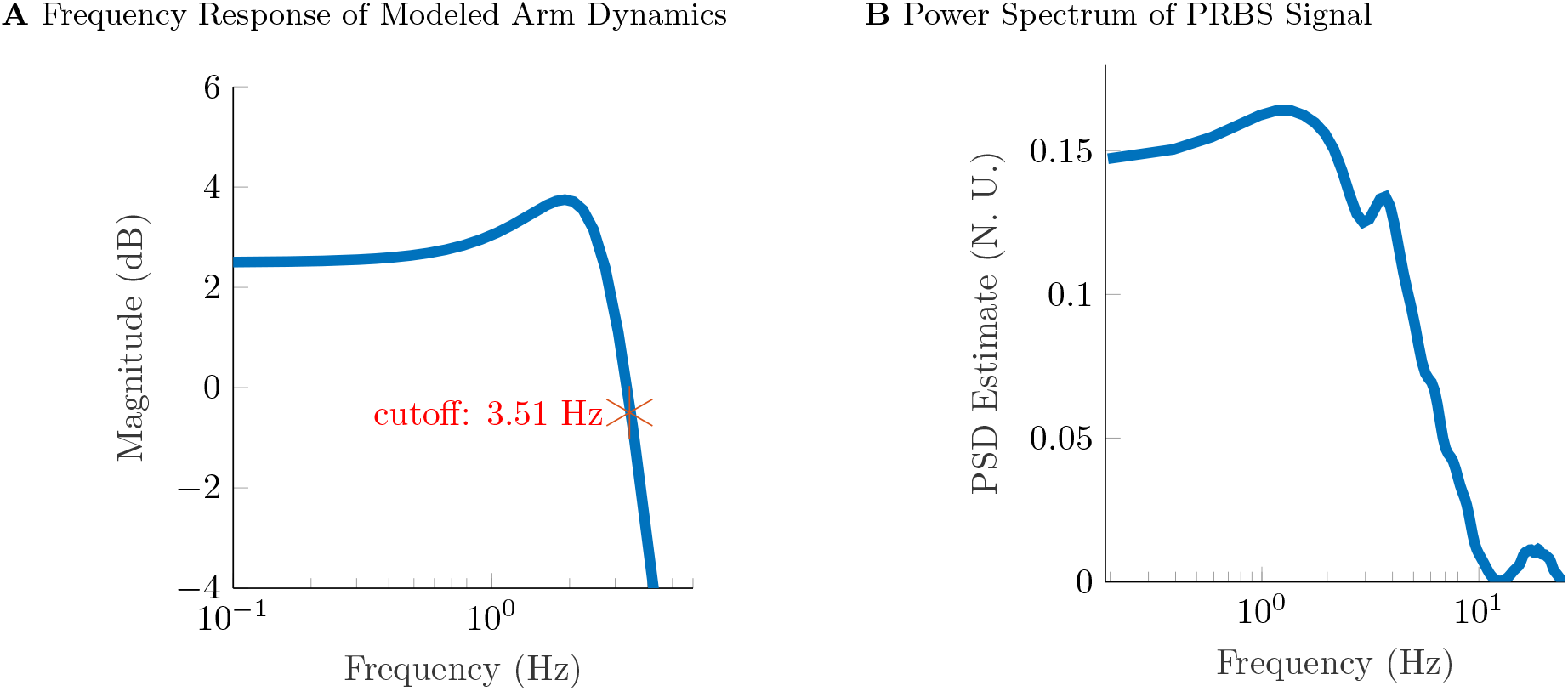
**A:** Bode plot of combined admittance control and approximated arm dynamics with the highest bandwidth. The transfer function used for the approximation is described in Appendix A. **B:** Power spectral density of PRBS perturbation signal that confirms the frequency content in the band of interest (0.15-10 Hz).

We employed a PRBS perturbation with zero mean and frequency content that excited the subject’s arm between 0.15-10 Hz for 20 s. We selected a bandwidth of 10 Hz to exceed the expected bandwidth of the arm motion (3.5 Hz) and capture the attenuation of the arm response at frequencies beyond 3.5 Hz. The PRBS signal used in the experiments is shown in Fig. 7. The power spectral density of the PRBS signal, shown in Figure 3B, confirms that the effective bandwidth of the perturbation was 0.15-10 Hz.

We heuristically determined the amplitude of the PRBS signal via experimentation to evoke angular displacements within the system’s physical limits. We used perturbations with an amplitude equal to +*/−* 1 *N · m*. The resulting standard deviation of the position signal was approximately 4 degrees for each perturbation epoch. The PRBS perturbations applied in the experiments were 20s long and were spaced between randomly selected 10 or 20 s rest periods when the subject’s arm was moved to the initial position. The same PRBS sequence was applied to all epochs across conditions (OFF and DBS) to compare parameter estimates associated with the same perturbation.

#### 2.5.4. Least-Squares Estimation of Arm Viscosity and Stiffness

We identified the inertial, viscous, and stiffness parameters of the arm using least squares estimation [14]. To estimate these parameters, the acceleration 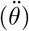 and position (*θ*) of the arm were reconstructed from the velocity commanded to the actuator. We used the velocity commands instead of the velocity measurements for parameter estimation to minimize the effect of sporadic communication jitters caused by our asynchronous communication between Simulink Desktop Real-Time and the actuator. Because the actuator’s torque and bandwidth were higher than those needed to generate the PRBS trajectories, the commanded velocity was an accurate approximation of the actual velocity. We used a forward finite difference time derivative of the velocity to calculate the arm acceleration, which is given by 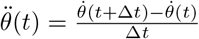, where Δ*t* is the sample time. The position was found through a forward finite difference time integral given by 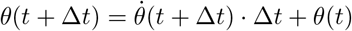.with *θ*(0) = 0.

Forces voluntarily applied by the subject to the robotic system can lead to low-frequency drifts in the position signal as a result of integrating the acceleration produced by the measured torque twice to obtain the position signal. To minimize the effect of voluntary motion on the estimation accuracy, we suppressed low-frequency drifts in the position signal using a Butterworth high pass filter with a cutoff frequency of 0.20 Hz.

We used Equation 3 to define the relationship between known data and unknown parameters in the least-squares estimation method. This relationship is given by

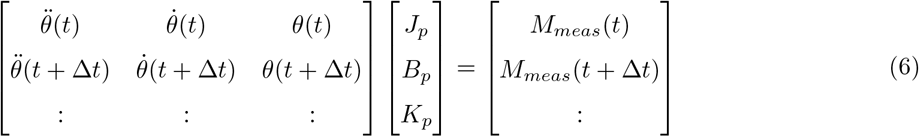

which has the form *Ax* = *b* with *x* = [*J*_*p*_ *B*_*p*_ *K*_*p*_]^*T*^. The least squares solution to *x* is 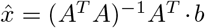 *· b*. We estimated inertia, viscosity, and stiffness parameters for each epoch associated with a PRBS sequence. Epochs in which the arm reached the position limits of the robot were excluded from the analysis. Epochs were also excluded if the goodness of fit statistic, *R*_*squared*_, was below zero.

The combined inertia of the robot and subject’s arm (*J*_*p*_) should not vary across conditions (OFF, DBS). Variations in the arm motion across these conditions can be attributed to changes in viscous damping and stiffness associated with DBS therapy. Consequently, we estimated the inertia *J*_*p*_ using the approach described in the previous subsection and then calculated the viscosity and stiffness parameters with a given constant inertia. The inertia was set equal to the median value of the inertia estimates across all epochs and experimental conditions recorded. After finding the median inertia estimate, the viscous damping and stiffness parameters were estimated using a least-squares minimization applied to the following expression.

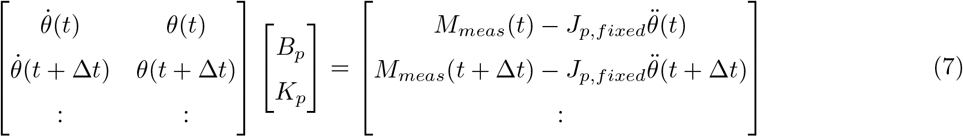

### 2.6. Data Collection and Statistics

Independent observations of the impulse and work scores were calculated based on epochs (data segments) of 20 s length associated with a PRBS sequence to assess statistical differences between observations across conditions (OFF, DBS). We tested the two experimental conditions over nine non-consecutive days. The subject’s rigidity was evaluated in the OFF condition each day with DBS either following or preceding the OFF condition. To minimize a possible sustained (wash-out) effect of DBS on rigidity, we conducted assessments in the OFF condition 15 min after turning DBS off. We evaluated DBS conditions for four days. Table 1 shows the number of epochs recorded to calculate the angular impulse score, work score, viscous damping, and stiffness.

**Table 1:** Number of epochs used for rigidity measurements in each condition across experiments.

| Rigidity Measurement | Condition | Total Epochs |
| --- | --- | --- |
| Corrected mUPDRS Score | OFF | 27 |
|  | DBS | 12 |
| Angular Impulse Score | OFF | 92 |
|  | DBS | 46 |
| Viscosity and Stiffness Estimation | OFF | 78 |
|  | DBS | 23 |

We made no assumptions about the distribution of each rigidity measurement. Therefore, a non-parametric, two-sided Wilcoxon rank-sum test was used to evaluate whether the medians of the viscous damping and stiffness in the DBS condition were equal to the median in the OFF condition (null hypothesis).

### 2.7. Simulation Environment

We created computer simulations of the robot motion to characterize how individual changes in viscous damping and stiffness affect the angular impulse and work scores and to validate the approach to estimating the arm’s damping and stiffness. Simulations without the admittance controller were created based on the differential equations of Equation 3 to characterize how changes in damping and stiffness altered impulse and work scores. We embedded the admittance controller in the simulation to validate that the least-squares estimation accurately predicted the damping and stiffness values when sensor noise was added to the simulated torque measurements. We modeled this noise as a white Gaussian process with zero mean and standard deviation equal to the standard deviation calculated from the torque transducer when no force or torque was applied to the robot. We used the same PRBS signals used in the experiments to implement the simulations and validate the least-squares estimation. The simulation was also used to assess how well our second-order linear model predicted the experimental data.

## 3. Results

### 3.1. Blind Rating Scale Scores

We corrected the mUPDRS scores for the individual examiner bias and variance with a linear mixed-effects (LME) model. The median and interquartile range of the corrected mUPDRS scores are shown Figure 4A. The LME model formula defined the experimental condition as a fixed effect and the individual examiners as a random effect on the mUPDRS scores. The F-test of the corrected DBS vs. OFF comparison indicated that the mUPDRS scores in the DBS condition were significantly lower than those in the OFF condition (p = 0.020).

**Figure 4:**
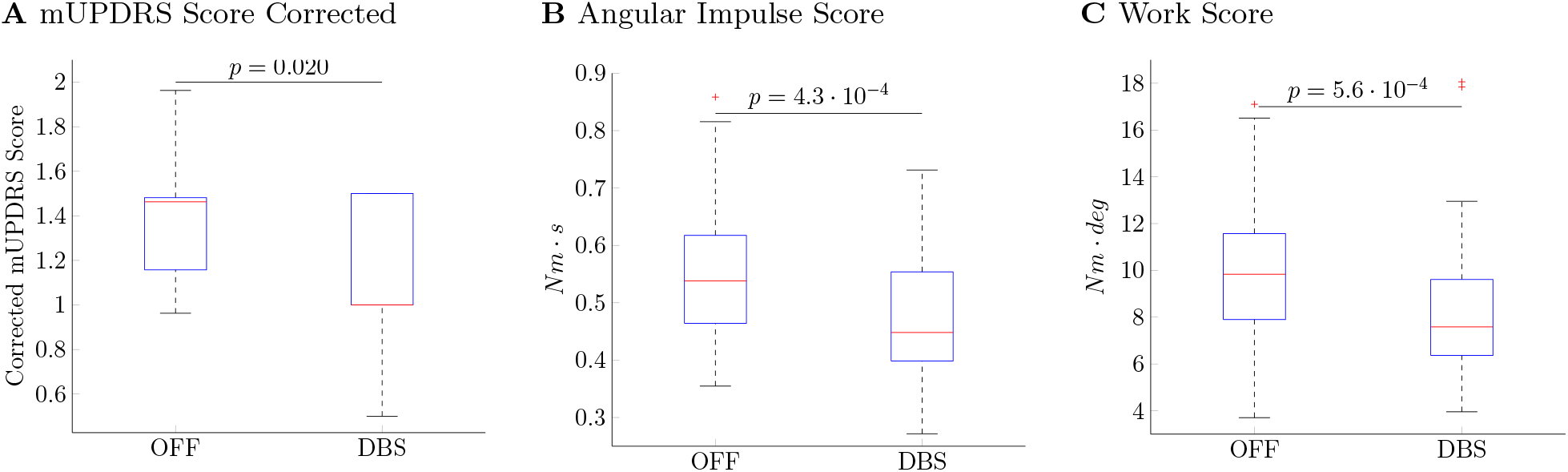
**A:** mUPDRS scores corrected using the linear mixed-effects (LME) model. The LME model corrects for the individual examiner bias and variance. **B:** Angular impulse scores for each condition. **C:** Work scores for each condition.

### 3.2. Simulation of Viscous and Stiffness Effects on Angular Impulse and Work Scores

We used the computer simulations of the robotic system and subject described in Section 2.7 to characterize how changes in viscous damping and stiffness parameters affect the impulse and work scores. The viscous damping and stiffness parameters used in the computer simulations were taken to be the median values estimated across all conditions (OFF, DBS) using the parameter estimation approach described in Section 2.5. We used these values to simulate scenarios with realistic viscous damping and stiffness. The median parameters were varied separately by 25%, 50%, 150%, and 175%, and then the change in work and angular scores relative to the baseline (median) values was calculated.

Figure 5A shows that an increase in the stiffness or viscosity results in an increase in the angular impulse score. Deviations in the angular impulse score due to stiffness changes are larger than those due to viscosity changes. This result indicates that the angular impulse score is more sensitive to changes in stiffness but cannot capture individual variations in the stiffness or viscosity alone. The work score is a monotonically increasing function of viscous damping but a monotonically decreasing function of the stiffness. See Figure 5B. Therefore, a simultaneous increase (or decrease) in both stiffness and viscous damping can lead to insignificant changes in the work score. This effect is primarily due to the sinusoidal nature of the perturbation together with the dependence of viscosity and stiffness on velocity and position, respectively.

**Figure 5:**
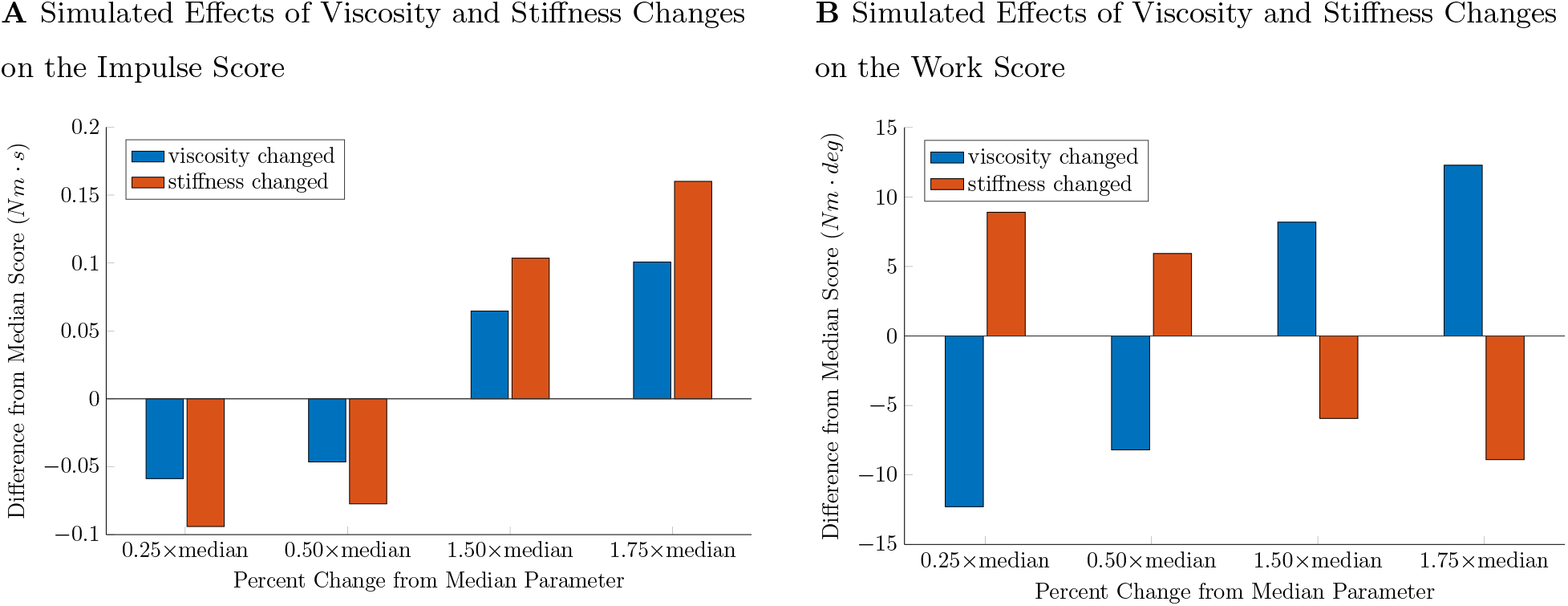
**A:** Changes in the impulse score following a percentage change in the viscous damping or stiffness parameter. The nominal inertia, viscous damping, and stiffness used in this analysis were equal to the median estimates of these parameters. The values of these parameters are (*Jp* = 0.004400, *Bp* = 0.1300, *Kp* = 1.130 (rad)). A 1.0 Hz sinusoidal perturbation to the position signal was used to create the total reaction torque in the computer simulations. **B:** Changes in the work score following a percentage change in the viscous damping or stiffness parameter.

### 3.3. Angular Impulse and Work Scores from Experimental Data

Sample traces of the torque as a function of the robotic system’s angular position trajectories are shown in Figure 6. The measured torque exhibited an asymmetric behavior when the elbow deflection was at its maximum flexion or extension. These asymmetries were likely generated by a combination of factors, including the subject’s alignment with the robot and the nonlinear relationship between the arm angular position and reaction torque.

**Figure 6:**
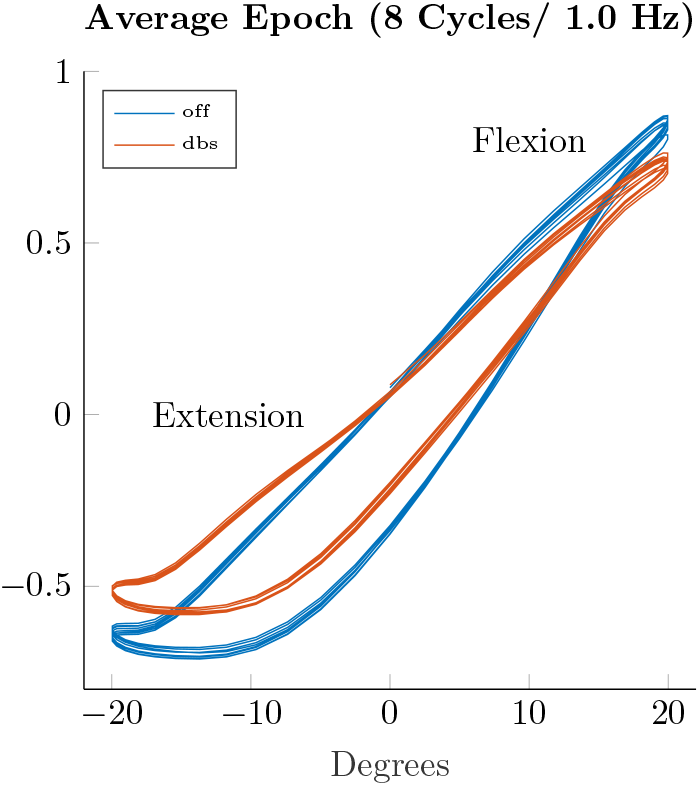
Torque vs. angular position trajectories when sinusoidal perturbation is applied to the position commanded to the robot.

According to the Wilcoxon rank-sum tests, the angular impulse scores in the DBS condition were significantly lower than those in the OFF condition (p = 4.3 *·* 10^*−*4^). Figure 4B shows the interquartile ranges and median of the angular impulse scores in the studied conditions. The work scores had a significant decrease from the OFF to the DBS condition (p = 5.6 *·* 10^*−*4^, Wilcoxon rank sum tests). See Figure 4C.

### 3.4. Simulation of Parameter Estimation Routine

Numerical simulations of the model estimation routine, with the same PRBS signal used experimentally and noise levels observed in the torque transducer, were carried out to characterize the estimation errors. In these simulations, we employed the moment of inertia (*J*_*P*_), viscous damping (*B*_*P*_), and stiffness (*K*_*P*_) values estimated from the experiments (*J*_*p*_ = 0.0045, *B*_*p*_ = 0.1368, *K*_*p*_ = 1.751). See experimental results in Section 3.5. A total of 100 simulations with distinct noise samples resulted in mean errors for *J*_*P*_, *B*_*P*_, and *K*_*P*_ equal to *−*1.9 *·* 10^*−*7^, *−*2.6 *·* 10^*−*7^, and *−*1.9 *·* 10^*−*7^, respectively. The standard deviation of these errors were 9.6 *·* 10^*−*7^, 1.05 *·* 10^*−*6^, and 1.4 *·* 10^*−*6^.

### 3.5. Parameter Estimation from Experimental Data

We evaluated the accuracy of the mathematical models estimated via least-squares by comparing the torque calculated using these models and the torque measured by the transducer during experiments. We calculated the torque signal in two different ways. First, we calculated the *estimated* torque signal as the estimated parameters (*J*_*p*_, *B*_*p*_, *K*_*p*_) multiplied by the angular acceleration, velocity, and position measurements, respectively. Second, we used the simulation environment described in subsection 2.7 to calculate the *simulated* acceleration, velocity, position, and torque directly from the input PRBS signal. The PRBS signal used in the experiments is shown in Figure 7A. Figure 7B illustrates how well the estimated (model) torque describes the torque measured during an experiment in the OFF condition (median values *J*_*p*_ = 0.004400, *B*_*p*_ = 0.1300, *K*_*p*_ = 1.310).

**Figure 7:**
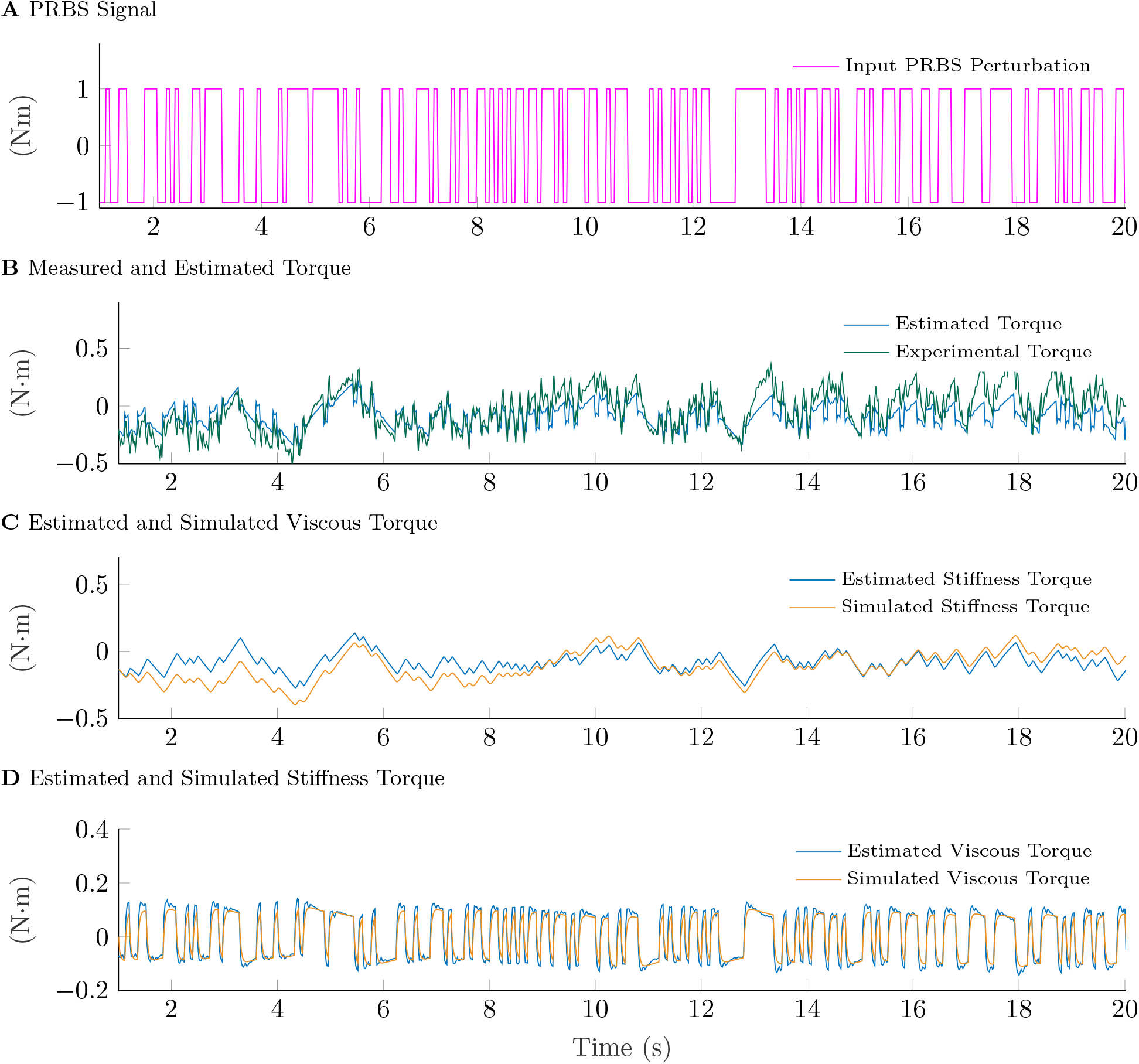
**A:** PRBS signal used during experiments. **B:** Torque measured during an experiment in the OFF condition together with the estimated torque, which was computed based on the estimated model parameters. The estimated torque was calculated as the inertia, viscosity, and stiffness estimates multiplied by acceleration, velocity, and position signals, respectively. **C and D:** Torque components associated with the stiffness and viscous damping. The simulated torque was calculated based on the acceleration, velocity and position signals derived via integration of the input PRBS perturbation.

Figures 7C and 7D illustrate the individual torque contributions from the position- and velocity-related torques (viscosity and stiffness) calculated using both the estimated and simulated torque. The similarity between the measured and simulated torque indicates that the PRBS signal drove the arm and robot’s motion. Therefore, in this experiment, the subject’s active resistance had a negligible effect on the arm and robot motion. The amplitude of the stiffness-related torque is larger than the viscosity-related torque at low frequencies, but smaller at high-frequencies. Discrepancies between the estimated and simulated stiffness are larger than those observed in the viscous torque. This difference occurs because the double integration of small drifts in the measured torque can lead to drifts in the position signal larger than those observed in the velocity signal.

Figure 8A shows the interquartile range and median of the inertia estimates across experiments for each condition. The Wilcoxon rank-sum test indicated that the inertia did not have statistically significant changes between conditions (p = 0.165 for OFF vs. DBS). Figures 8B and 8C show the median and interquartile ranges of the viscosity and stiffness estimates for each condition. Both the viscous damping and stiffness exhibited a significant decrease from the OFF to the DBS condition (p = 5.0 *·* 10^*−*4^ for viscous damping and p = 1.8 *·* 10^*−*4^ for stiffness).

**Figure 8:**
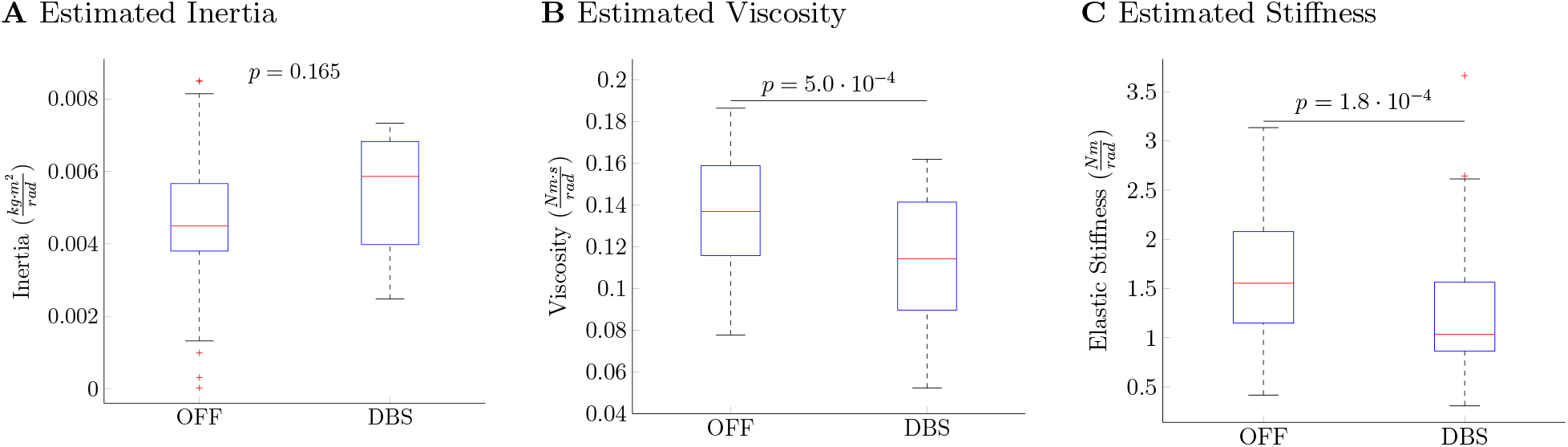
**A:** Median and interquartile ranges of the inertia estimates in each condition. These estimates were found through the least squares approach. No significant differences in the inertia estimates were found between conditions. **B:** Median and interquartile ranges of the viscous damping estimates for each condition. A fixed inertia of 0.0045 *kg · m*^2^ was used for the estimation. The viscosity in the DBS condition was smaller than in the OFF condition (p = 5.0 *·* 10^*−*4^, Wilcoxon rank sum test). **C:** Medians and interquartile ranges of the stiffness estimates in the OFF and DBS condition based on a fixed inertia. The viscosity in the DBS condition was smaller than in the OFF condition (p = 1.8 *·* 10^*−*4^, Wilcoxon rank sum test).

### 3.6. Effect Sizes of Condition Comparisons

Table 2 shows the effect sizes of each rigidity assessment for the OFF vs. DBS condition comparisons. We calculated these effect sizes from the r-value proposed by Cohen using the z statistic of the Wilcoxon rank-sum test [15].

**Table 2:** P-values and effect sizes from statistical comparisons of rigidity measurements between the ON-and OFF-DBS condition.

| Rigidity Measurement | Comparison | p-value | effect size (r) |
| --- | --- | --- | --- |
| Corrected mUPDRS Score | OFF vs DBS | 0.04 | 0.18 |
| Angular Impulse Score | OFF vs DBS | $4.3 \cdot 10^{-4}$ | 0.30 |
| Work Score | OFF vs DBS | $2.8 \cdot 10^{-4}$ | 0.31 |
| Viscosity | OFF vs DBS | $4.0 \cdot 10^{-4}$ | 0.35 |
| Stiffness | OFF vs DBS | $1.8 \cdot 10^{-3}$ | 0.37 |

Cohen’s guidelines state that effect sizes with large, medium, and small effects are generally around 0.5, 0.3, and 0.1, respectively. The p-values of the angular impulse score, work score, viscosity, and stiffness were calculated from the Wilcoxon rank-sum test. We calculated the mUPDRS p-values from the F-test of the linear mixed-effects model.

## 4. Discussion

We developed a systematic methodology to isolate the effect of velocity and angular position on rigidity and estimate the elbow’s viscosity and stiffness. Our methodology enables the subject being tested to freely move the arm while torque perturbations are injected to characterize the rotational arm dynamics. Our experimental data demonstrated the capability of model-based parameter estimation to estimate changes in viscous damping and stiffness when comparing the OFF and DBS conditions in an animal model of PD. These data indicate that the proposed parameter estimation approach can help researchers and clinicians quantify treatment effectiveness and understand the physiological mechanisms underlying the manifestation of both stiffness and viscosity in PD. We also compared the proposed methodology with rigidity assessment approaches typically used for research, including the mUPDRS scale and the angular impulse and work scores.

Unlike the mUPDRS scale, the angular impulse and work scores remove human biases from quantifying rigidity. These scores provide a holistic representation of all the different factors contributing to the subject’s rigidity, including the reaction moments that are a function of the angular position, velocity, and acceleration. The comprehensive nature of the angular impulse and work scores is their strength and weakness. Capturing all of the subject’s dynamics with one metric hides the individual factors that may contribute to the subject’s rigidity.

All rigidity assessment methods evaluated in this study (mUPDRS, angular impulse, work, parameter estimation) revealed that DBS resulted in lower scores than the OFF condition. The estimates of viscosity and stiffness had the largest effect sizes among all other rigidity scores. A larger effect size translates to a more substantial difference in the distributions, which suggests that for the studied subject, our parameter estimation is better at differentiating between the DBS and OFF conditions than the work, impulse, and corrected mUPDRS scores.

While the mUPDRS scores aggregated across examiners via the LME model suggest that rigidity was decreased from the OFF to the DBS condition (p=0.02), the effect size of this difference is significantly smaller than the effect sizes associated with the objective quantification methods. The coarse-scale (low-resolution) of the mUPDRS scores and implicit biases in perception within and across evaluators are factors that might have influenced the small effect size in the mUPDRS assessments.

Although the experimental data suggests that the work score can differentiate changes in rigidity, our computer simulations revealed that this score is a monotonically increasing function of the viscous damping for the parameter ranges studied here but a monotonically decreasing function of the stiffness. These observations suggest that the work score is not suitable to evaluate rigidity in parkinsonian subjects because a simultaneous increase or decrease in viscous damping and stiffness can result in underestimating overall changes in rigidity.

The proposed parameter estimation method was based on a second-order, linear, time-invariant approximation of the coupled arm and robot dynamics. The actual dynamics of the arm and robot may exhibit high-order terms and nonlinearities that our simplified model does not capture. Nevertheless, the accuracy of the second-order linear model is remarkable given its simplicity (see Fig. 7) and suggests that this model is of sufficient complexity to study stiffness and viscosity. When estimating the arm dynamics, the inertia values for each condition did not differ statistically, validating our assumption that the subject and robot’s inertia is constant across experiments. Variance in the estimated inertia, viscosity, and stiffness can be attributed to active resistance from the subject, deviations in the robot-subject alignment, and high-order and nonlinear dynamics not captured by our second order linear model of the arm-robot motion.

Our experimental results indicate that viscosity and stiffness in the DBS condition of the studied animal were both significantly lower than in the OFF condition. These results support the hypotheses that 1) parkinsonism is associated with changes in both viscous damping and stiffness, and 2) DBS reduces these two rigidity components. Our data also suggest that model-based parameter estimation is suitable for assessing the therapeutic effect of DBS and other treatments on viscosity and stiffness in PD. Future studies can leverage the model-based estimation approach presented here to characterize how descending inputs from the brain alter muscle groups’ organization and how this alteration relates to changes in viscosity and stiffness in PD and during DBS. One can also use the robotic system and methodology described herein to develop automated programming routines for the selection of DBS parameters (pulse-width, frequency, and current amplitude).

### 4.1. Limitations

A significant challenge for measuring rigidity in nonhuman animals is the active resistance affecting any assessment approach. Although we attempted to identify the arm dynamics associated with a PRBS perturbation injected into the torque signal while allowing the subject to freely move, we acknowledge that active resistance occurring at the same frequency of the PRBS signal can play a role in the variability of the parameter estimates. Using electromyography (EMG) recordings could help detect epochs with active movement and isolate these epochs for analysis..

The experimental results that we present in this study are based on a single subject. They are evidence that the proposed model-based estimation approach can be used to quantify changes in rigidity in a large animal model of PD. However, generalizations about the effects of DBS on stiffness, viscosity, angular impulse scores, and work scores require a study with a larger cohort of subjects.

## Acknowledgments

We thank Maya Stoller, Devyn Bauer, and Ethan Marshall for performing the mUPDRS scores as well as Professors Matthew Johnson, Tim Kowalewski and Colum MacKinnon for their insightful feedback through-out this study.

## Appendix A. Estimation of Cutoff Frequency

We determined the bandwidth (cutoff frequency) of the arm rotational dynamics from a second-order model of the subject’s arm. The forearm mass was approximated by a human body-forearm weight ratio of 1.5% [16, 17] with the subject’s body mass at 7.7 kg (animal weight). The forearm inertia was approximated as 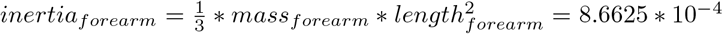, which is the inertia of a rod rotating about it’s end [18]. The robot arm’s rotational inertia was calculated via computer-aided design (CAD) software and was found to be equal to 4 *·* 10^*−*3^ *kg · m*^2^. We determined the total system inertia from the combined subject, robot, and tuned admittance controller inertia (0.0008*kg · m*^2^*/rad*). The total inertia was equal to 0.0052*kg · m*^2^*/rad*.

The subject’s stiffness was approximated as 1.5 Nm/rad from studies of elbow elastic muscle properties in humans with and without Parkinson’s disease [19, 20]. The robot’s shaft and arm were considered rigid bodies. Therefore, the stiffness properties of the robot were considered negligible. Because the controller does not modulate stiffness, the total system stiffness was equal to the approximated subject’s stiffness.

We approximated the effective viscous damping of the subject’s elbow as 0.06*Nm · s/rad* according to viscosity ranges found in a study of viscosity properties of the human elbow [21]. The tuned controller viscous damping (0.028*Nm · s/rad*) was added to the subject’s approximated viscosity for a final system viscosity of 0.0088*Nm · s/rad*.

To estimate the maximum frequency cutoff of the system, we accounted for 50% error in the approximated inertia, viscosity, and stiffness. We added this uncertainty to account for potential inaccuracies in the approximation from literature values and differences between rhesus macaque and human anatomy. Then we identified the transfer function with the largest bandwidth given the parameter ranges within the uncertainty bounds. The transfer function with the largest cutoff frequency, described in equation A.1, was used to create the bode plot of Figure 3A.

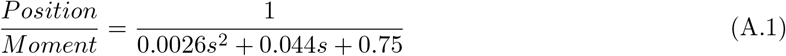

